# The Curious Case of the Golden Orb – Relict of *Relicanthus daphneae* (Cnidaria, Anthozoa, Hexacorallia), a deep sea anemone

**DOI:** 10.64898/2026.04.17.719276

**Authors:** Steven R. Auscavitch, Abigail Reft, Adena B. Collens, Christopher Mah, Merlin Best, Charlotte Benedict, Estefanía Rodríguez, Marymegan Daly, Allen G. Collins

## Abstract

The discovery and collection of the enigmatic Golden Orb by the NOAA Ship *Okeanos Explorer* and ROV *Deep Discover* in deep Alaskan waters during 2023 has yielded substantial interest by the scientific and public communities alike. Initial field identifications of the specimen collected at 3,250 meters depth ranged from an egg mass to sponge to microbial biofilm. Here we characterize the biology and ecology of the Golden Orb, as well as other specimens of similar appearance identified since the collection of the original material. Through an integrative taxonomic approach including morphological analysis and genomic characterization of the Golden Orb, we identified the presence of cnidocytes of the spirocyst type (restricted to Hexacorallia), as well as metazoan DNA, from which we were able to derive complete mitochondrial genomes and Ultra Conserved Elements. These results indicate that the Golden Orb and a similar specimen from deep equatorial waters represent remnant cuticles belonging to the geographically widespread deep-sea anemone ally *Relicanthus daphneae*. We also document the presence of cuticle from a collected specimen of *R. daphneae* from the Southern Ocean and *in situ* photographic evidence of similar cuticles beneath living individuals. These findings underscore the extent to which the biodiversity and organismal biology of obscure deep sea fauna broadly remain unresolved and highlight the value of whole-specimen collections and rigorous taxonomic follow-up in telepresence-enabled ocean exploration.

## Introduction

The mission of NOAA Ocean Exploration (NOAA-OE) is to explore and document unknown regions of the ocean, with a focus on US waters deeper than 200 meters within its Exclusive Economic Zone (EEZ) for national benefit. As part of the effort to achieve this goal, NOAA-OE operates the NOAA Ship *Okeanos Explorer*, from which it deploys cutting edge technology to achieve its mission. Using multibeam sonar systems, well over 2 million square kilometers have been mapped with NOAA Ship *Okeanos Explorer* (*Okeanos Explorer*), including significant portions of the Pacific (Albano et al. 2025), as well as Earth’s largest deep water coral reef (Hoy et al., 2025). The ROV *Deep Discoverer* (D2, deployed from *Okeanos Explorer*) allows for livestreaming with public and shore-side scientists to follow along on expeditions that provide critical observations on deep marine ecosystems (Cuellar et al. 2024). High-definition video has allowed for the documentation of many novel biological behaviors, associations and interactions (e.g., Wicksten et al., 2017; Sigwart et al., 2019; Mah 2022; Vecchione 2019; Pratt et al., 2023). In addition, since 2015, *Okeanos Explorer* has been equipped to collect specimens that become part of the collections of the Smithsonian National Museum of Natural History (NMNH), where they are cataloged and preserved, fostering their study into perpetuity. For example, 40 specimens collected via *Okeanos Explorer* have been designated holotypes and 15 as paratypes to support the description of 44 species and seven genera (Reft et al., 2026).

During the course of *Okeanos Explorer* expeditions, it is not uncommon that encountered organisms are not immediately recognized. In most of these instances the appropriate specialist or needed information is simply unavailable for on-site determination, but follow-up investigation with additional references and the broader community quickly resolves these uncertainties. Data from *Okeanos Explorer* has shown that imagery can provide important and powerful data for our understanding of the biology and ecology of deep-sea organisms (e.g., Auscavitch et al., 2020, Quattrini & Demopoulos 2016).

However, sometimes real mysteries exist and imagery alone only raises questions. Such is the case of the Golden Orb. During a 2023 expedition in the Gulf of Alaska, as part of the Seascape Alaska Campaign, scientists encountered a golden, mound-shaped object attached to a rock and with an apparent torn opening on its surface (Fig. 1). Because of the distinctive appearance of the specimen and its mysterious nature, the Golden Orb attracted considerable speculation and significant public interest, resulting in numerous independently written accounts (e.g., *USA Today*, *Forbes*, *Miami Herald*, *Smithsonian Magazine*). Fortunately, the specimen was collected using a suction sampler and sent to NMNH where its morphology and genetics could be examined.Through an integrative approach examining morphology and genetics of the specimen (USNM_IZ_1699903), we have determined that the Golden Orb is the organic remnant of *Relicanthus daphneae* (Daly, 2006), a deep-sea anemone.

**Figure 1:**
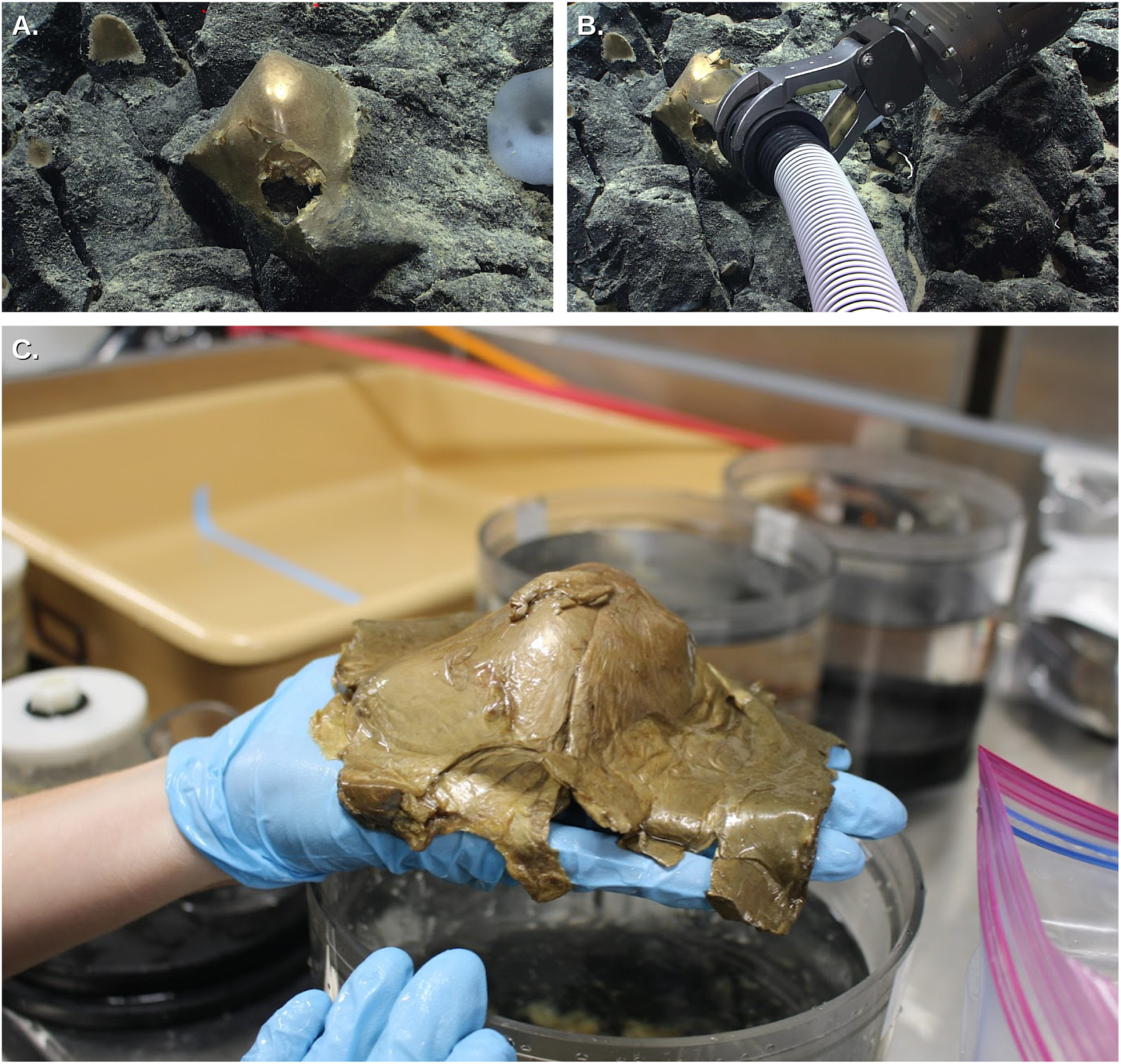
The‘Golden Orb’ specimen collected by NOAA Ship *Okeanos Explorer* during expedition EX 23-06 collected at a depth of 3,250 m. A. *In situ* photo before collection. B. Photo during collection by suction sampler. C. Specimen out of water in the wet lab. Photos credits: NOAA Ocean Exploration.

## Materials & Methods

### Golden Orb specimen preparation and Identification of other materials

The Golden Orb specimen ( USNM 1699903; Fig. 1) was collected on August 30, 2023 during Dive07 of the Seascape Alaska 5 Expedition, which was deployed to explore an unnamed volcanic feature southwest of Walker Seamount in the Gulf of Alaska (55.013, -140.833). The specimen was found among a field of small glass sponges (family Euplectellidae Gray, 1867) growing on exposed basalt cobbles at a depth of 3,250.76 m and recovered whole using a suction sampler. Video of the collection can be viewed on <u>NOAA Ocean Exploration’s website</u>. Immediately following its recovery the Golden Orb was imaged and preserved in 95% ethanol at room temperature. An additional subsample of tissue (Biorepository Number AR0OK12) was preserved in cryogenic tubes, also in 95% ethanol, and at room temperature for later archiving at the NMNH Biorepository.

During an 2021 Schmidt Ocean Institute expedition of the US Pacific Islands Heritage Marine National Monument (Discovering Deep-sea Corals of the Phoenix Islands 2) with the R/V Falkor and ROV SuBastian, an earlier example of Golden Orb-like material was identified from a sample collected at 2,939 m at an unnamed seamount south of Titov Seamount within the US EEZ (USNM_IZ_1770968). Encountered on a rocky surface of a large boulder, the specimen was originally thought to be sponge or microbial biofilm (Fig. 2A). An approximately 10 cm specimen was collected with the ROV manipulator by gently scraping the rock surface and suctioning up the fragment (Fig. 2B). Later examined in the ship’s lab, the specimen was noted as flakey and embedded with small foraminifera tests that composed common deep-sea sediments at this site. The entire fragment was preserved in 95% ethanol and held at room temperature.

**Figure 2:**
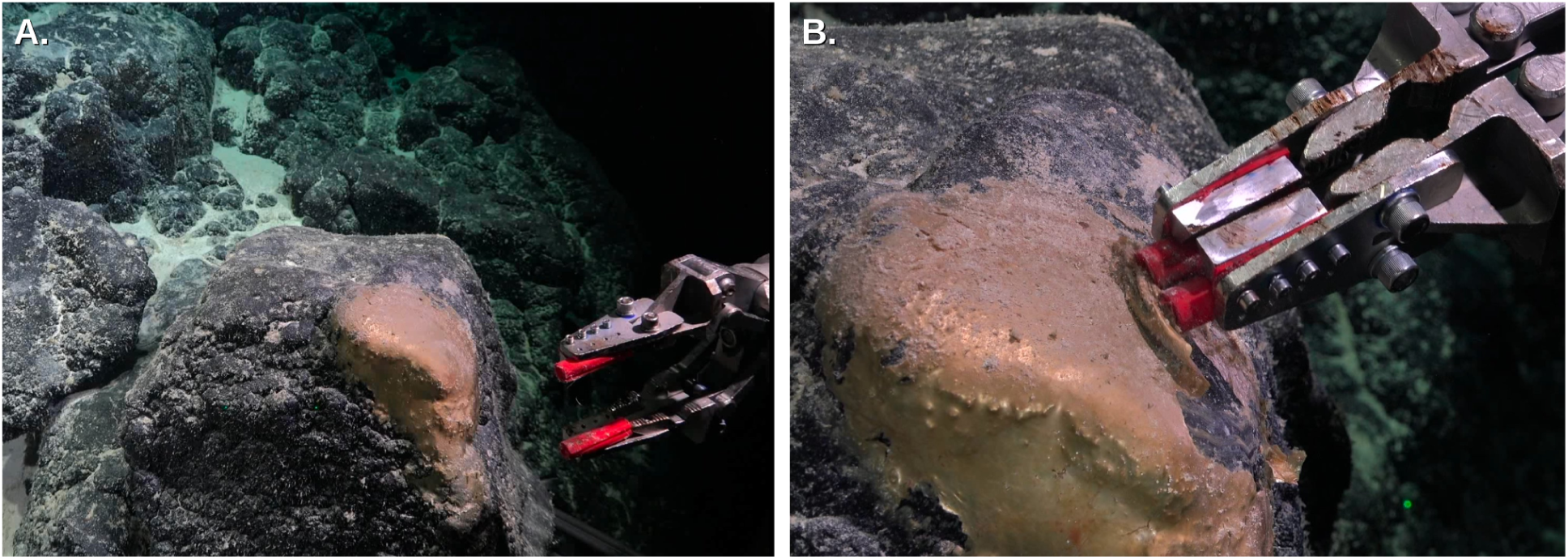
Seafloor imagery of Rock Skin material from specimen FK21-449 (USNM_IZ_1770968) from R/V Falkor cruise FK210605 in the US EEZ surrounding Howland and Baker Islands. (A) Size and (B) zoomed image indicating the laminated texture and soft consistency of the structure.

### Morphological characterization with photography, light microscopy and scanning electron microscopy

Whole specimen images of USNM 1699903 (Fig. 3) were taken and stacked using Helicon Focus (version 8.3.11, Helicon Soft Ltd, http://www.heliconsoft.com). USNM 1770968 is thin and flat and did not require stacking. Small sections of multiple regions of both samples were removed, squashed on a slide and imaged on Olympus BX63F DIC compound microscope. Spirocysts were measured from photographs using ImageJ. Sections from both the inner and outer parts of USNM 1699903 were removed, critical point dried, mounted, and sputter coated for scanning electron microscopy (SEM). These samples were imaged using a Zeiss EVO MA15 SEM. For comparative purposes, we also examined a partially dissected whole specimen of *Relicanthus daphneae* (to be deposited at the NHMUK) collected near hydrothermal vents in the Scotia Ridge (Southern Ocean) in 2012 at 3,948 m.

**Figure 3:**
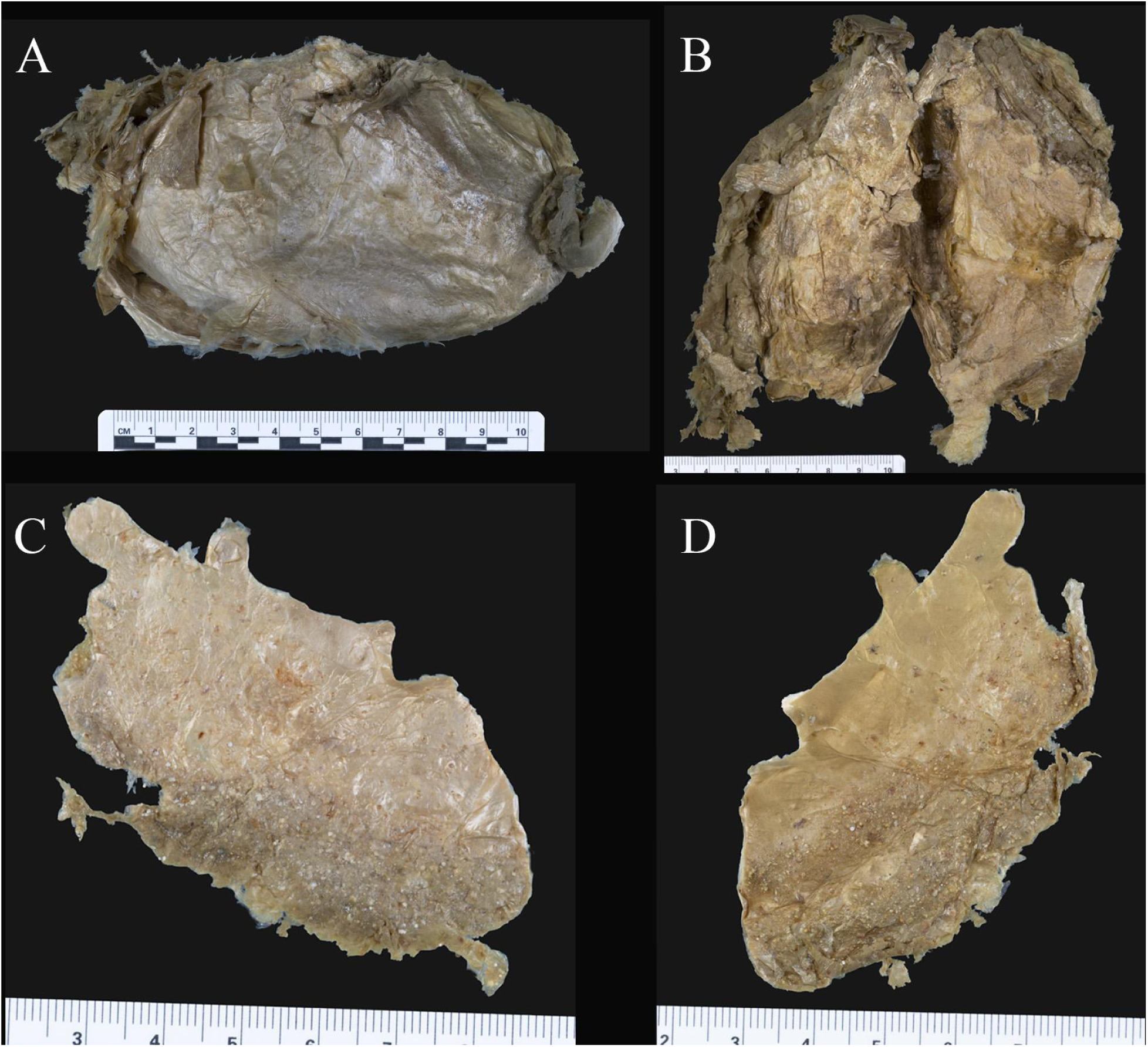
Whole specimen images of the Okeanos specimen USNM_IZ_1699903 (A-B) and Falkor specimen FK21-449 (USNM_IZ_1770968) (C-D). A. External surface of USNM_IZ_1699903. B. Internal surface of USNM_IZ_1699903. C. External surface of USNM_IZ_1770968. D. Internal surface of USNM_IZ_1770968.

### Molecular Characterization

All DNA extracts were isolated using the AutoGenPrep 965 automated DNA extraction system (AutoGen, Holliston, MA, USA) according to the manufacturer’s tissue protocols. After PCR using Folmer COI primers failed on an initial DNA extraction, several tissues were subsampled from the Golden Orb specimen including the top and underside of surface, as well as internal tissue (Table 1). From specimen FK21-449 (USNM_IZ_1770968), owing to its thinness (Fig. 3), two DNA extractions were made from surface-only tissues, i.e., top and bottom of surface (Table 1). Libraries were enzymatically sheared and prepared with the NEB Ultra II FS DNA library preparation kit for Illumina (New England Biolabs), aiming for an insert size of around 400 bp. Library amplification was performed with ten cycles of PCR, following the manufacturer’s recommended chemistry and thermocycler settings. We used iTru y-yoke adapter stubs and iTru unique dual indices instead of NEB adapters and indices, adjusting the adapter quantity based on DNA concentrations per NEB’s guidelines. We used Quant-ITdsDNA HS assay (Thermo Fisher Scientific, Waltham, MA, USA) and High Sensitivity D1000 ScreenTape (Agilent, Santa Clara, CA, USA) to determine library size. Equimolar pooled libraries were sequenced with 150 bp paired-end reads on a NovaSeq X Plus (Illumina Inc., San Diego, CA, USA).

**Table 1.** Tissues sampled in this study (NCBI BioSamples SAMN56737688-694) and genomic data obtained (NCBI SRA SRR37877116-122) and products generated (NCBI Nucleotides PZ234282-83) from sequencing via Genome Skimming.

| **<u>Sample Label</u>** | **<u>Reads Before Filtering</u>** | **<u>Reads After Filtering</u>** | **<u>mtGenome Derived</u>** | **<u>Coverage mtGenome</u>** |
| --- | --- | --- | --- | --- |
| USNM_IZ_1699903a_Internal | 19,480,702 | 19,191,670 | no | na |
| USNM_IZ_1699903b_Internal | 54,760,524 | 53,917,980 | no | na |
| USNM_IZ_1699903_Internal-deep | 39,629,484 | 39,146,534 | no | na |
| USNM_IZ_1699903_Underside-surface | 128,267,796 | 126,996,356 | yes | 83.54 |
| USNM_IZ_1699903_Surface | 157,285,824 | 155,761,512 | yes | 118.56 |
| FK21_449_Surface | 73,831,454 | 73,026,662 | yes | 19.52 |
| FK21_449_Underside-surface | 131,617,682 | 130,027,916 | yes | 8.98 |

Reads from each library were processed to remove adapters and low-quality sequences using FastP (Chen et al., 2018) and assembled with metaSPAdes (Nurk et al., 2017). Mitofinder (Allio et al., 2020) was used to identify mitochondrial genomes with draft annotations of gene regions. Derived mitochondrial genomes were aligned with a reference mitochondrial genome from *Relicanthus daphneae* (MK947129). After read pre-processing, full-length SSU rRNAs were assembled with phyloFlash (Gruber-Vodicka et al., 2020). In the phyloFlash workflow, reads were mapped to reference SSU rRNA sequences in the SILVA 138.2 database (Quast et al., 2013; Yilmaz et al., 2014). Only reads which mapped to the reference SSU rRNAs were assembled de-novo with SPAdes (Bankevich et al., 2012). Taxonomic identity of the SSU rRNAs was characterized by the closest reference in the SILVA database and verified using BLASTn to NCBI (Altschul et al., 1990; Supplementary Table S1).

Assembled contigs were searched using the PHYLUCE pipeline and matched conserved element loci to the *hexacoral-v2* bait set (25,288 baits targeting 2,476 loci, 1,127 UCE and 1,349 exon loci) (Cowman et al., 2020) using phyluce_assembly_match_contigs_to_probes with min-coverage of 70% and min-identity of 70%. Final identified loci were extracted using phyluce_assembly_get_match_counts, aligned with MAFFT (Katoh et al., 2002) using phyluce_align_seqcap_align, and trimmed internally using Gblocks with default parameters. The taxon occupancy matrix was obtained for 50% completeness. Phylogenetic analyses including 164 taxa (Supplementary Table S2) were completed using IQTree v2.4.0 (Nguyen et al., 2015, Minh et al., 2020). The best-fit substitution model was selected using ModelFinder based on the Bayesian Information Criterion (BIC). The resulting topology was rooted with representatives of Octocorallia.

Metagenomes were assembled, binned and quality scored using the nf-core/mag pipeline v. 3.0.2 implemented in Nextflow (Krakau et al., 2022). Raw reads for each sample were pre-processed by trimming adapters and for quality with fastp (Chen et al., 2018), then assembled using MEGAHIT (Li et al., 2016). Then, samples were co-assembled by similarity of SSU read groupings. Then, MetaBAT2 was used to bin contigs by nucleotide frequencies and co-abundance patterns by the sample group (Kang et al., 2019). Metagenome-assembled genomes (MAGs) were evaluated using bacterial single-copy marker genes from the GTDB bac120 set. For each bin, we recorded the number of bac120 markers detected as single-copy genes, duplicated genes, and missing genes, and used these counts as a proxy for CheckM-style completeness and contamination. Bins with ≥108 of 120 markers present as single-copy genes (≈≥90% completeness) and low marker duplication were classified as high-quality (HQ) MAGs, whereas those with 60–107 markers (≈50–90% completeness) were classified as medium-quality (MQ) and retained for downstream analyses; all other bins were considered low-quality and excluded. Taxonomic assignments were obtained from the GTDB-Tk bacterial workflow, which places MAGs in a reference-based phylogenetic tree and resolves taxonomy from domain to genus (and species where possible) based on marker-gene phylogeny, ANI, and RED-based criteria for novelty.

## Results

Initial examination of the gross morphology of USNM_IZ_1699903 revealed the specimen to have no indication of typical animal anatomy (mouth, gut, muscle tissues, etc.), but rather to consist of a loose aggregation of fibrous material covered by a smooth, layered surface (Fig. 3). Similarly, additional specimens (FK21-449; USNM_IZ_1770968) displayed similar characteristics, i.e., somewhat golden and reflective, that was observed and collected in 2021 at 2,952 m during Schmidt Ocean Institute’s R/V Falkor cruise FK210605 in the US EEZ surrounding Howland and Baker Islands in the central Pacific (Figs. 2,3). Examination under light microscopy revealed the surface of both specimens to be packed with cnidocysts of the spirocyst form (Fig. 4), measuring an average of 73.95 µm in length and 6.06 µm in width (n=40). Spirocysts are an agglutinant cnidocyst type restricted to members of the cnidarian class Hexacorallia (Daly et al., 2003; Daly et al., 2007). The spirocysts documented in the column of *Relicanthus daphneae* range in length from 48.8-121.4 µm, and in width from 2.9-5.5 µm (Daly 2006). The lengths of the orb spirocysts are well within this range, though the widths were often wider. However, most of the orb spirocysts were partially discharged which has been demonstrated to have a negligible effect on length, but a significant effect on width, the discharged cnidae being significantly wider than undischarged (Ryland & Lancaster 2004). Therefore, the spirocysts found in both orb samples are consistent with those found in *R. daphneae*.

**Figure 4:**
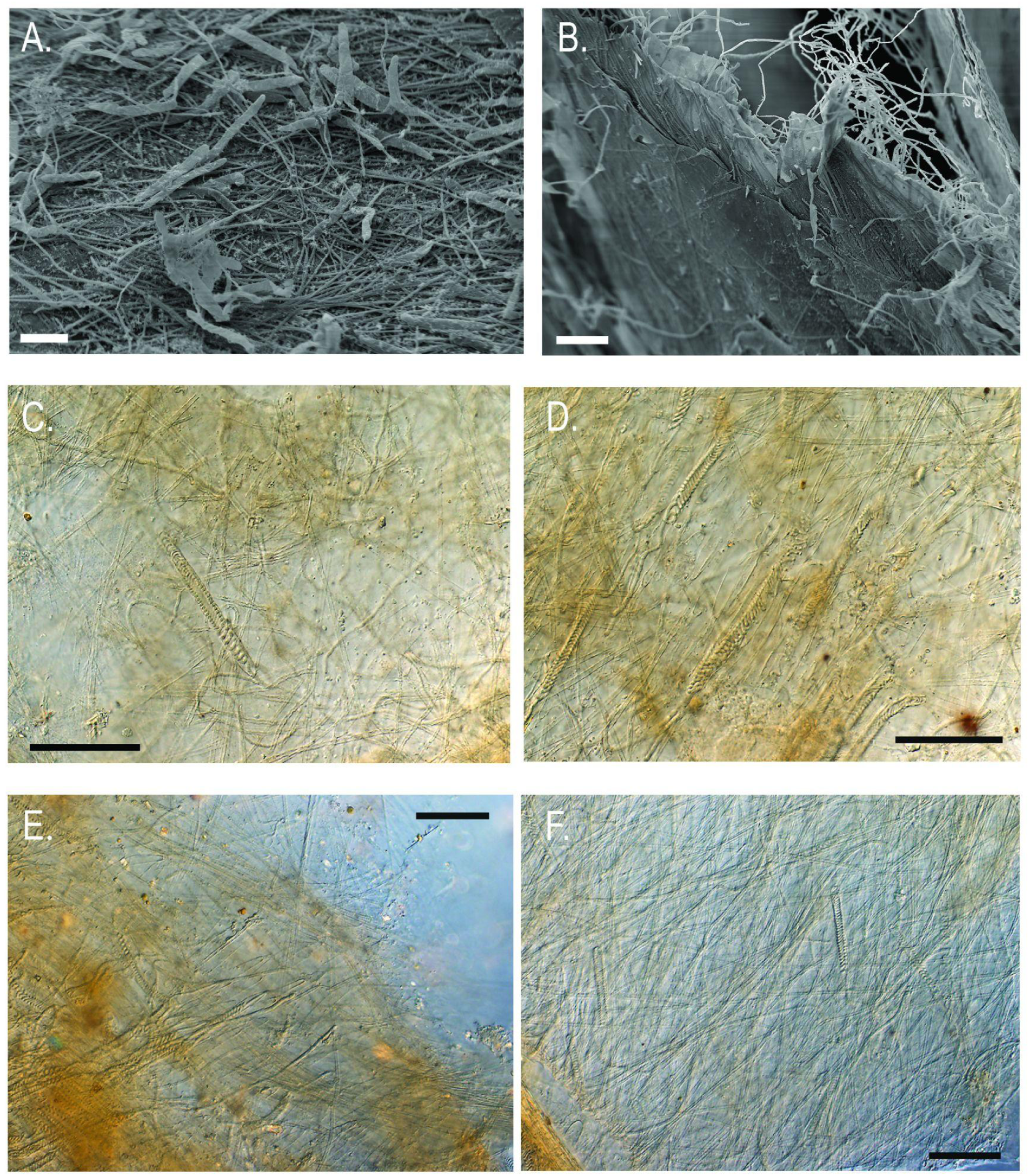
Microscope images of samples. A.-B. SEM images of Okeanos specimen USNM_IZ_1699903, C.-F. Light microscope images, scale bar:50 µm. The layers of the inner and outer surfaces consist of spirocyst capsules (fired and unfired) and their tubules. A. Image of the inner surface of USNM 1699903. Scale bar: 30 µm. B. Image of the outer surface of USNM 1699903, scale bar: 40 µm. C.-D. Images of layers from USNM_IZ_1699903. E. Image of a layer of the inner surface of USNM 1770968. F. Image of a layer of the outer surface of USNM 1770968.

Initial DNA barcoding of the mitochondrial COI gene using Folmer primers did not yield conclusive sequence data for any recognizable organism, likely due to co-amplification. Deeper whole genome sequencing was conducted on five subsamples of USNM_IZ_1699903 and two subsamples of the Falkor specimen FK21-449 (USNM_IZ_1770968). Sequencing yielded datasets ranging between approximately 19 to 155 million reads per sample (Table 1). PhyloFlash analyses indicated that all samples contained small subunit of the ribosome (SSU rRNA) of *Relicanthus*, but with varying proportions relative to SSU from other organisms (Fig. 5). The SSU of additional organisms from all domains of life were assembled by PhyloFlash (Gruber-Vodicka et al., 2020). A single representative SSU of ammonia-oxidizing *Nitrosopumilus* sp. Archaea was assembled from all but one of the datasets, whereas multiple and diverse bacterial representatives were recovered in all datasets. We also detected a handful of SSUs from non-metazoan taxa (various rhizarians, a possible chytrid fungus, and a discoban). In addition, several other metazoan taxa, including one nemertean, one nematode and one octocoral in USNM_IZ_169903; with low coverage other than the nemertean (22X). Both samples of FK21_449 yielded SSU of an octocoral (Supplementary Table S1).

**Figure 5:**
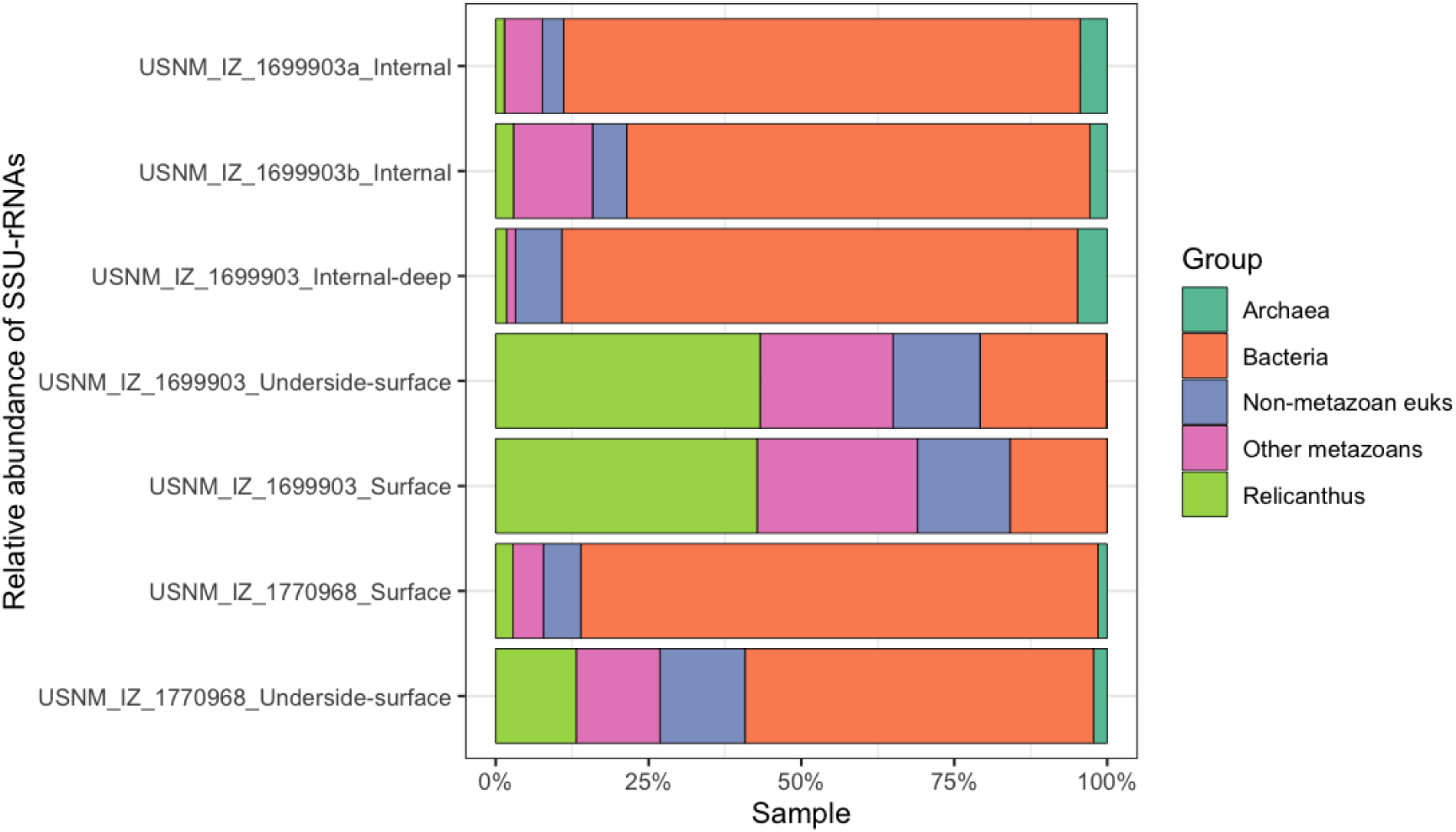
Relative abundance of SSU reads that map to *Relicanthus*, other metazoans, non-metazoan eukaryotes, Bacteria, and Archaea in each of our analyzed samples (Table 1).

Complete, circularized mitochondrial genomes, 17,724 bp in length, were obtained from four samples (two each from vouchers USNM_IZ_1699903 and FK21-449; USNM_IZ_1770968), with read coverage ranging from roughly 9X to 118X (Table 1). The replicate samples from each voucher were identical to one another, but the two vouchers differed by 17 bp (99.9% identical). The reference mitochondrial genome from *Relicanthus daphneae* (MK947129) is 17,727 bp long and differs from the USNM_IZ_1699903 and FK21-449 genomes at 29 and 16 positions, respectively. Overall the pairwise divergences among the three mitochondrial genomes are 0.096%, 0.138% and 0.065%. Alignment revealed that the three base pair length difference was due to three N’s (among other ambiguities) in the reference genome that appeared as insertions when compared to the mitochondrial genomes derived from our material.

UCE loci from the hexacoral-v2 bait set were identified in samples USNM_IZ_1699903_underside-surface, USNM_IZ_1770968_Surface, and USNM_IZ_1770968_Underside-surface, with the most (n=205) being found in USNM_IZ_1699903_Surface. In phylogenetic analyses including this sample, Ceriantharia was revealed as the sister group to the remaining hexacorals, among which Actiniaria was the earliest diverging lineage. USNM_IZ_1699903_Surface branched with another exemplar of *Relicanthus daphneae* and the two formed a clade sister to (Antipatharia(Corallimorpharia+Scleractinia)).

Application of bac120 marker-based thresholds yielded a set of medium- and high-quality MAGs from our samples across internal and surface samples (Supplementary Table S3). High-quality MAGs included a deeply novel Gammaproteobacteria genome (1699903_Internal-deep.1.fa) affiliated with order WLWR01 and a nearly complete Alphaproteobacteria genome (Surface.22.fa) within Rhizobiales, family Rhizobiaceae. Both bins exhibited >95% bac120 marker alignment coverage and marker profiles consistent with ∼90–98% completeness and low duplication. Additional medium- to high-quality MAGs were recovered from Gemmatimonadota (Internal-deep.8.fa; Longimicrobiales, UBA6960), Chloroflexota (Surface.1.fa; Dehalococcoidia, UBA2979/UBA6627), Bacteroidota (Surface.10.fa; Flavobacteriaceae, CANLXL01), Verrucomicrobiota (Surface.12.fa; Chthoniobacterales, CAMASH01), and Planctomycetota (Surface.17.fa; Gimesia), all of which showed ≥85% marker MSA coverage, indicating near-complete genomes suitable for comparative and functional inference.

Metagenome-assembled genomes (MAGs) recovered from our samples revealed a community that is taxonomically consistent with deep-sea benthic microbiomes, while also exhibiting substantial phylogenetic novelty. Across both bacterial and archaeal MAGs, we identified representatives of Thermoproteota (*Nitrosopumilus*-like AOA), Pseudomonadota (Woeseiales, Nitrosomonadaceae, diverse Alphaproteobacteria), Chloroflexota (Anaerolineae), Acidobacteriota, Gemmatimonadota, Planctomycetota, Sumerlaeota, and several candidate phyla. Many of these MAGs were classified only at the level of uncultured genera or families and exhibited sub-radius average nucleotide identity (ANI) values relative to their closest references, indicating that the *Relicanthus* cuticle niche harbors numerous undescribed species and potentially deeper lineages.

## Discussion

The curiosity surrounding the Golden Orb has yielded considerable excitement among deep-sea scientists, exploration enthusiasts, and the media alike. To date, a cursory survey of international news media, broadcast media, and topical science and nature outlets has generated 27 unique articles about the Golden Orb according to Google search. At the Smithsonian Institution, inquiries to the status of the Golden Orb persist to the present day with frequent public inquiries since its collection in September 2023. Through an integrative taxonomic approach we conclude that the Golden Orb is not an object of alien origin, but rather one of Earth’s own inner space which still holds many secrets left to uncover. In addition, the Golden Orb is not an egg of some animal or a sponge. Instead it represents a remnant of cuticle and tissue belonging to the enigmatic deep-sea anemone, *Relicanthus daphneae*, which has been inhabited by diverse microscopic life forms. While these findings provide closure to the case of the Golden Orb, the curiosity of the specimen, and speculation surrounding its discovery, underscores the importance of exploration and scientific collections.

*Relicanthus daphneae* has a fascinating evolutionary history. It was originally described as a species within the actiniarian genus *Boloceroides* Carlgren, 1899 (see Daly 2006), but the specimen collection that led to its description happened 30 years after its initial discovery, highlighting the lag time associated with taxonomy. Since the first explorations of chemosynthetic environments, *R. daphneae* was recognized as a commonly observed inhabitant of vent periphery habitats on the East Pacific Rise, but it was initially thought to be a species of ceriantharian (Segonzac & Doumenc in Desbruyeres & Segonzac, 1997). Later phylogenetic analyses based on five genes suggested that it did not fall within Actiniaria or any of the other orders in the cnidarian class Hexacorallia, leading to the creation of a new genus, family, and suborder for this animal (Rodríguez et al., 2014; Xiao et al., 2019). Analyses of complete mitochondrial genome sequences provided moderate support for the hypothesis that *R. daphneae* was sister to the remaining Actiniaria (Xiao et al., 2019), but these analyses also suggested that sponges were derived from within the cnidarian subphylum Anthozoa, a result with other phylogenetic hypotheses, prompting additional inquiry. Phylogenomic analyses failed to recover a close relationship between *R. daphneae* and Actiniaria, instead favoring, albeit without full support, the species as forming the sister group to a clade consisting of the orders Antipatharia, Corallimorpharia and Scleractinia (McFadden et al., 2021; Quattrini et al., 2023). A more recent investigation of the phylogeny of Actiniaria included *R. daphneae*, where it formed the sister group to Actiniaria, but most other hexacoral taxa were not included in this analysis (Benedict et al., 2024). Our phylogenetic analysis of UCEs concurs with other results based on this type of data, finding *Relicanthus* to form the sister group of a clade uniting Antipatharia, Corallimorpharia, and Scleractinia, with that entire clade sister to Actiniaria (Fig 6.).

**Figure 6:**
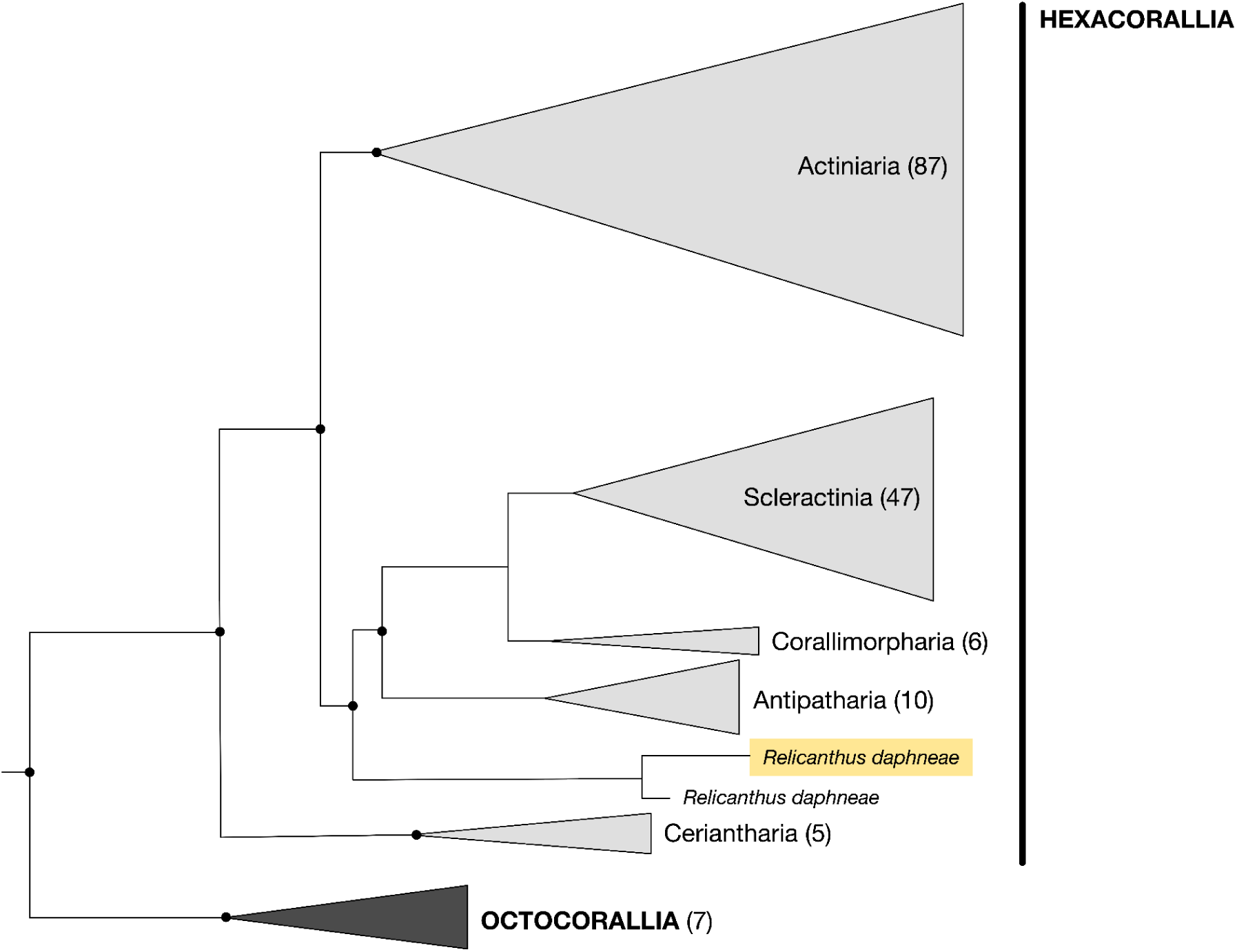
Maximum likelihood phylogeny of Anthozoa estimated from a UCE dataset (94 loci) using a 50% occupancy matrix. Sequences were aligned with MAFFT and filtered using Gblocks prior to phylogenetic analyses in IQ-TREE. Best-fit substitution model was selected using ModelFinder based on the Bayesian Information Criterion (BIC). Nodes with ultrafast bootstrap support ≥50 are indicated by dark circles. The Golden Orb subsample USNM_IZ_1699903_Surface is highlighted in yellow. The full topology with all terminal taxa can be found in the supplemental material (Supplementary Figure S1).

Compared to many anthozoans, *Relicanthus* are relatively easy to identify from video footage and as specimens. *Relicanthus* can have a polyp diameter of approximately 0.3 m, which makes it among the largest-bodied polyps in Anthozoa. The smooth, cylindrical column bears a dense crown of very long, sinuous tentacles, which may obscure the oral disc and mouth. The tentacles are generally the same purple, pinkish, or red as the column, but may be darker or lighter depending on the relative state of contraction. Tentacles can be twice as long as the column is wide and vary in length, but are always strongly tapering and often coiled at the tips. When the animal is disturbed, tentacles may detach from the polyp; small muscles at the base of each tentacle allow them to pinch off or autotomize from the body. A contracted animal can be less than a quarter of the size of the expanded polyp, forming a low mound.

The original description of *R. daphneae* did not mention cuticle, and indeed there is none visible on the pedal disc of the type specimen (see Figure 2C, Daly 2006). However, one whole specimen collected at the periphery of hydrothermal vents in the East Scotia Ridge (Scotia Sea, Southern Ocean) was retrieved with several pieces of a loose, thin cuticle plate (Fig. 7). This cuticle is thin but multilayered (at least two layers of cuticle together, one of these layers is golden) and showed linear impressions corresponding to the mesenterial insertions of the pedal disc supporting that it was attached to the pedal disc at some point in the specimen’s life; we also found remains of the whole animal tissue still attached in a few spots of the broken cuticle (Fig. 7). Observations of animals *in situ* suggest that cuticle is left behind as the animal moves (Fig. 8B,D), suggesting that the animal can detach from it, which might explain its absence from many collected specimens.

**Figure 7.**
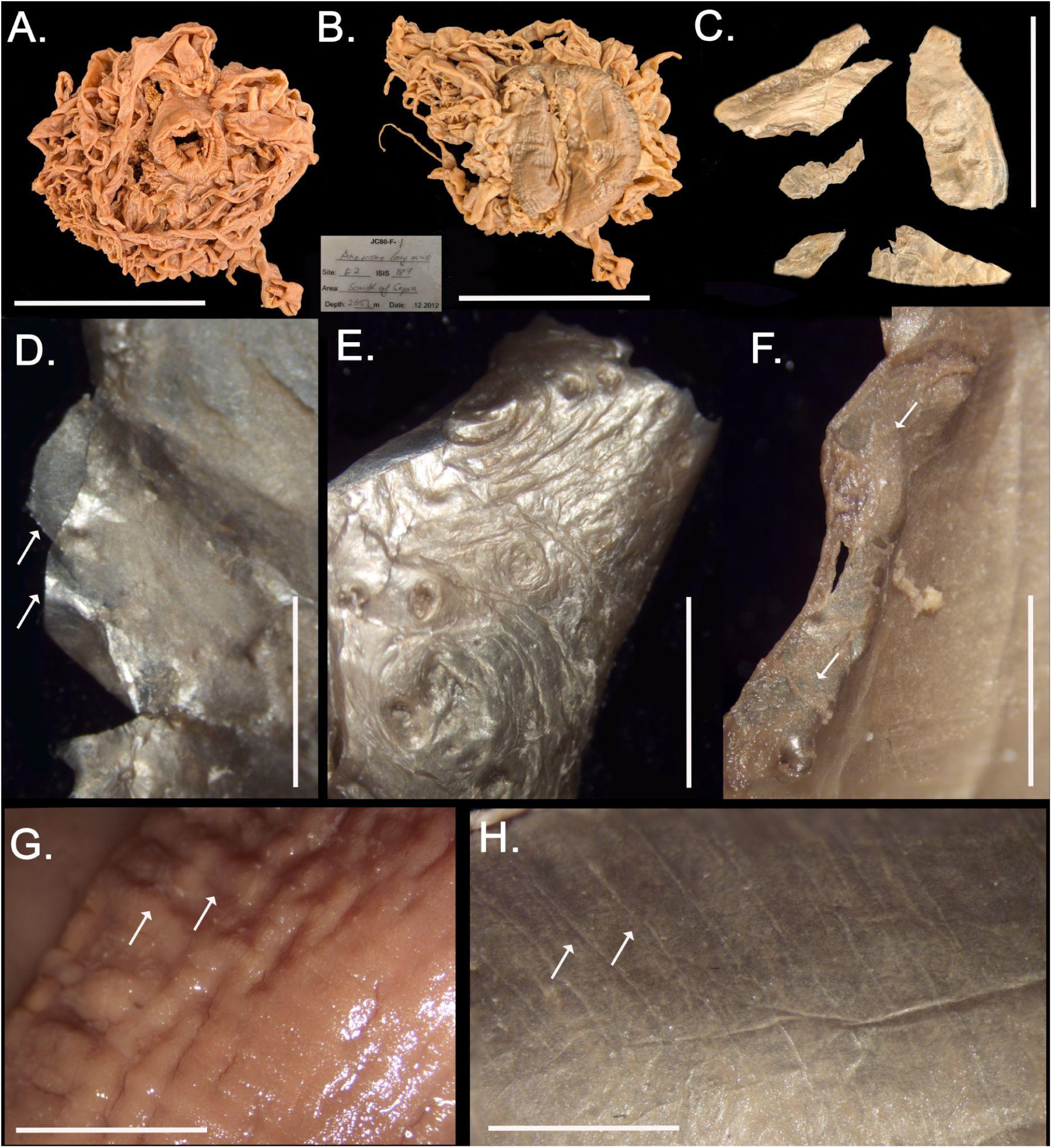
Specimen and cuticle remains of *Relicanthus daphneae* from the Scotia Ridge. A. & B. Oral and pedal disc view of preserved specimen. C. view of the cuticle remains. D. - F. Detailed view of the cuticle remains, note bilayered nature of cuticle (D.) and specimen tissue remains in the cuticle (F.). G. Close-up view of the pedal disc of the specimen, note indentations of the mesenterial insertions in the pedal disc (arrows). H. Details of cuticle remains mirroring marks of the indentations of the mesenterial insertions in the pedal disc. Scale bars: A.-B.: 7 cm; C.:5 cm; D.-F.: 1 cm; G.: 2 cm; H.: 1 cm.

**Figure 8:**
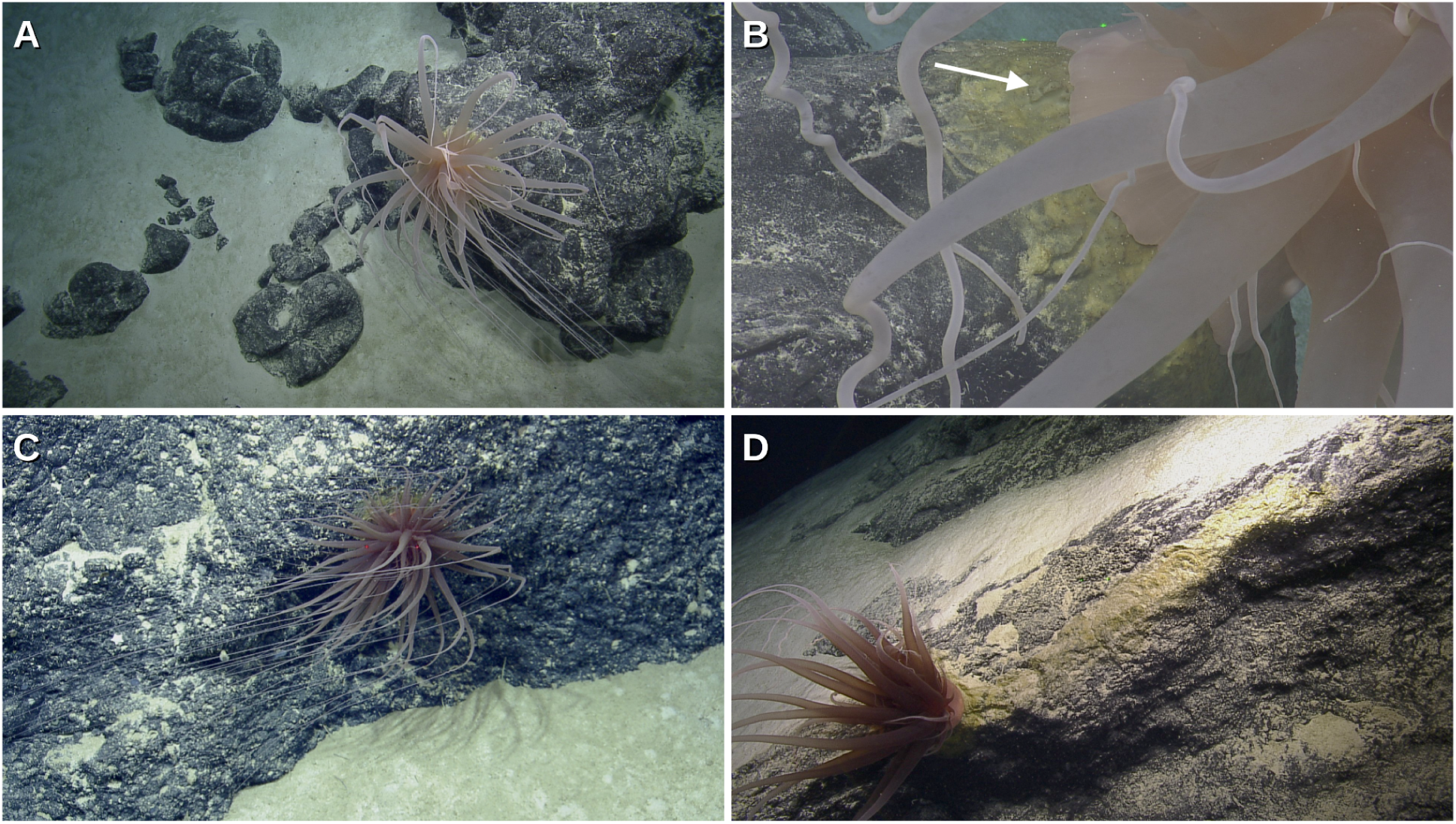
Photo records of *Relicanthus daphneae* associated with golden cuticle secreted by the pedal disc. The distance between red or green scaling lasers is 10 cm. A. A large individual from the central Pacific observed during E/V *Nautilus* expedition NA137 Kingman Reef & Palmyra Atoll exploration. B. Close-up of the same individual in A. showing detail of pedal disc secretions (white arrow). C. Another individual with cuticle observed during E/V *Nautilus* Expedition NA035 in the Anegada Passage, Caribbean Sea. D. Individual with trailing golden cuticle indicating persistence of *Relicanthus* cuticle layers as the individual moves across rocky surfaces. E/V *Nautilus* expedition NA137 Exploration of Kingman Reef & Palmyra Atoll. Photo credits: Ocean Exploration Trust.

Chitin synthase genes appear to be broadly distributed across Cnidaria, including in animals and specific life history stages where chitinous structures are unknown (Vandepas et al., 2023). Cuticle is most well characterized in the pedal disc/base from anemones that are symbiotic with hermit crabs (e.g., Dunn et al., 1980, Crowther et al., 2011, Gusmão et al., 2019 Yoshikawa et al., 2022). It is also well-documented in non-symbiotic species like those in the family Hormathiidae Carlgren, 1932 (e.g., *Hormathia* Gosse, 1859, *Phelliactis* Simon, 1892, etc.), which often have cuticle in the column that can be patchy and most obvious in the columnar tubercles. In species in the genus *Galatheanthemum* Carlgren, 1956 (G. *profundale* and *G. hadale* Carlgren, 1956, see Cairns et al., 2007), *Chitinactis marmara* Gusmão & Rodríguez, 2021, or *Octineon* Moseley in Fowler, 1894, the cuticle is unusually well developed and covers the column, or even forms a tube into which the anemone retracts (e.g., *G. profundale*). In most of these, the cuticle is primarily composed of chitin (Dunn & Liberman 1983; Yoshikawa et al., 2019) secreted by the epidermis. Most species of anemones that produce a well developed columnar cuticle inhabit deep waters (e.g. *Galatheanthemum*, *C. marmara*, *Octineon*); however, a correlation between cuticle and deep sea environments has not been demonstrated.

Observations of *Relicanthus* individuals are not altogether rare in the deep-sea, owing in part to the increased effort in deep-sea exploration in recent decades and its broadcast through online channels (e.g., oceanexplorer.noaa.gov/livestreams/, NautilusLive.org, or YouTube livestreams). However, the tendency to collect individuals of *Relicanthus* for scientific research is infrequent, likely because of their enormous size, and long, trailing tentacles that make them difficult to obtain through manipulator arm or suction sampling without substantial damage to the specimen. As a result, specimen records are fairly limited. Nevertheless, *R. daphneae* appears to be globally distributed and inhabits depths ranging from 1,667 to 3,948 meters (Fig. 9; Supplementary Table S4). The Global Biodiversity Information Facility (GBIF) records 32 occurrences of *R. daphneae*. However, we noticed that six of these records surprisingly had a shallow depth, and when interrogated stem from erroneously identified eDNA metabarcode sequences published by Obst et al. (2020). In fact, BLAST comparison of the ASVs in question showed unequivocally that the sequences were not from any cnidarian. This highlights the need to minimize false positive eDNA metabarcode identifications before they are shared with resources such as GBIF or the Ocean Biodiversity Information System (OBIS), in which we found the same errors. To the 26 confirmed records in GBIF, we can add our two records, as well as a recent observation from the Indian Ocean (Neufeld et al., 2024).

**Figure 9:**
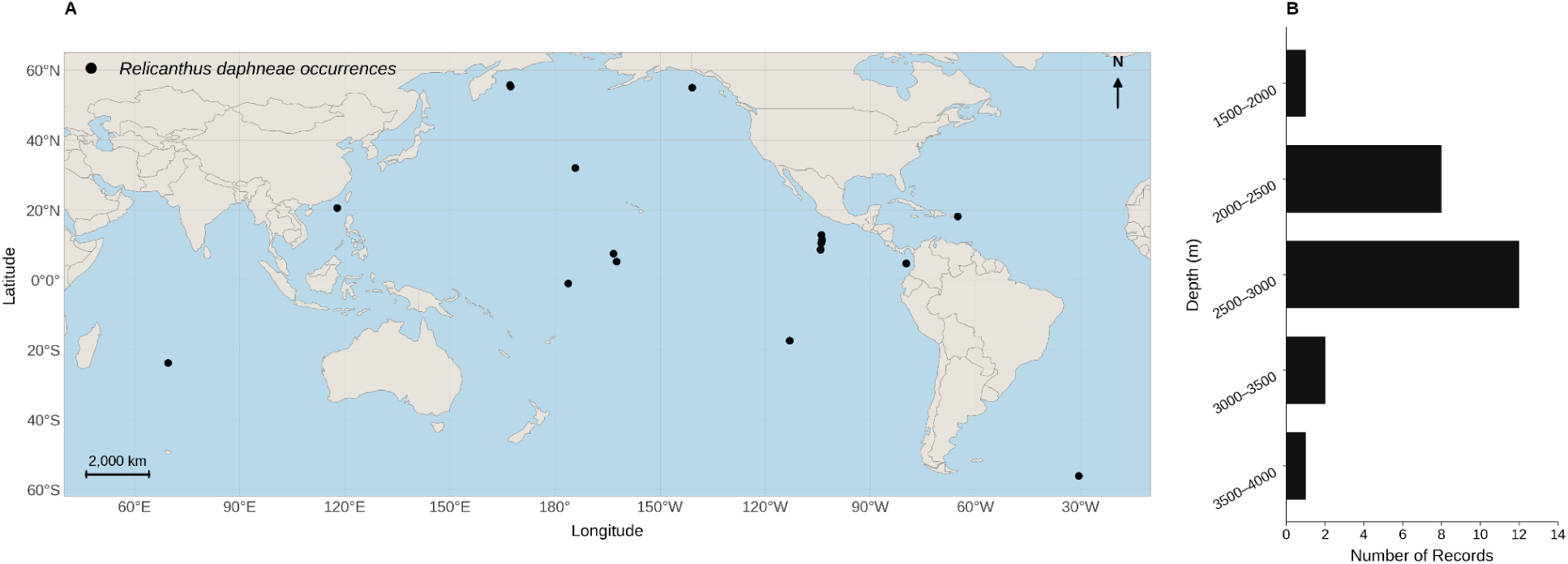
Global distribution of published records of *Relicanthus*. A. *Relicanthus daphneae* occurrences (n= 28) from video observations and specimen collections, including those referenced in GBIF, but excluding shallow-water eDNA-based occurrences that we interpret to be erroneous (see text). B. Bathymetric distribution bar chart indicating the range of depths of these occurrences where depth data were available.

Although genetic and morphological data confirm the identification of the taxon in question, explanation of the golden orb morphology remains a vexing issue. One possible interpretation is that the orb is a remnant of incomplete asexual reproduction. Some sea anemones are capable of pedal laceration, whereby the base of the polyp is abandoned and the upper portion of the animal moves away, leaving a stump of the body that then regrows a new polyp (Cary 1911). This is observed in anemones in the family Aiptasiidae Carlgren, 1924 and related taxa (e.g. Grajales & Rodríguez 2014). Partially regrown anemones that reproduce in this manner have an appearance somewhat similar to that of the Golden Orb, with a mound-like shape topped by an uneven convexity. In conjunction with the evidence for movement, this could explain one manner in which *Relicanthus* undergoes reproduction.

Because *R. daphneae* is known from few specimens and lives in such an inaccessible habitat, understanding of its biology largely comes from inference rather than direct observation. Many reports and observations are of multiple polyps in relatively close proximity; other anthozoans achieve this through asexual reproduction through e.g., pedal laceration, low dispersal of sexually produced offspring, or recruitment of unrelated larvae to a site through shared cues (Glon et al., 2020). The eggs of *R. daphneae* are large and very yolky; in other anthozoans, large, yolky eggs are characteristic of long-lived, dispersive offspring and so would argue against low dispersal. There is no evidence of anatomical irregularities in any specimens examined to date that would be expected of asexual reproduction through longitudinal fission, but asexual reproduction through regeneration of autotomized tentacles is likely, based on sea anemones like *Boloceroides* and *Bunodeopsis* Andres, 1881 (see Pearse 2002).

Video and photo records of *Relicanthus in situ* provide some insights to their biology and ecology. *Relicanthus* individuals tend to be encountered on, or perhaps settle preferentially, on the upper or current-exposed side of boulders which allows space to stream its long tentacles in the flow (Fig. 8A,C); specimens have been also found attached on the stalks of glass sponges. From these perched positions, they are able to capture prey, with a recent observation of an individual adjacent to a hydrothermal vent field capturing vent shrimp (*Rimicaris kairei* Watabe & Hashimoto, 2002; Neufeld et al., 2024). In revisiting high-quality imagery taken of *Relicanthus* collection events, we found that it is often possible to see cuticle secretions on which the live animal rests (Fig. 8B,D), although these are generally not visible when the animal is *in situ* because they do not extend beyond the margin of the pedal disc. Settlement among anemones is not necessarily fixed for life (Riemann-Zürneck 1998). Shallow-water anemones can creep along substrates (Bedgood et al. 2020), and deep deep sea anemones have been observed to employ various modes of locomotion including burrowing (Durden et al. 2015), rolling (Marchiò et al. 2025), passive drifting (Simon-Lledó et al. 2023, Dayton 1995), and active swimming (Ellis et al. 1969). Anemones in the genus *Boloceroides*, to which *R. daphneae* was originally assigned (Daly 2006), can actively swim in shallower waters (Josephson & March 1966), using their tentacles to row the body. This raises the possibility that *Relicanthus* may be able to move across the deep-sea landscape by evacuating its cuticle up or down slope with local changes in e.g., flow conditions, food availability, or to escape predation. Indeed, one image appears to show a trail of golden material indicating accumulation of secreted cuticle layers as an animal moved laterally across a rocky surface (Fig. 8D).

Metagenomes recovered from our samples reveal microbial assemblages with a pronounced nitrifier signature embedded in a diverse heterotrophic background. We recovered high-quality MAGs assigned to *Nitrosopumilus*-like ammonia-oxidizing archaea and Nitrosomonadaceae ammonia-oxidizing bacteria, alongside multiple Woeseiales, Chloroflexota (Anaerolineae), Acidobacteriota, Planctomycetota, Gemmatimonadota, and Sumerlaeota genomes. Taxonomically, these MAGs cluster with deep-sea sediment and seafloor biofilm lineages (Shi et al., 2023), many of which show substantial phylogenetic divergence from cultured representatives (Supplementary Table S3). The co-occurrence of archaeal and bacterial nitrifiers with heterotrophs typically associated with organic-rich sediments (Hoshino et al., 2020) points to active coupling of organic matter degradation and nitrogen transformations at the site of decaying *Relicanthus* tissue.

The strong representation of both archaeal and bacterial ammonia oxidizers suggests that the cuticle of a dead *Relicanthus* may act as a localized nitrification hotspot in the deep sea. As host-derived proteins and other nitrogen-rich biomolecules may be remineralized by heterotrophic taxa such as Woeseiales, Chloroflexota, and Acidobacteriota, the resulting ammonium pool appears to be rapidly oxidized by *Nitrosopumilus* and Nitrosomonadaceae. This tight spatial coupling between organic matter degradation and nitrification transforms the decaying cuticle into a micro-scale biogeochemical reactor, potentially influencing local redox conditions and nitrogen speciation on and around hard substrates.

## Conclusion

The deep sea represents a massive portion of the habitable space on Earth, and remains largely unexplored. That said, centuries of study have steadily built knowledge of deep-sea biodiversity, including the identities of new species and novel associations. However, it is still not unusual for exploratory surveys to present video observations and/or objects that defy immediate explanation. In some instances, the puzzle can be as simple as What is that? Here, we have resolved this basic question prompted by one such mysterious object. The Golden Orb is the organic relict of an animal that likely died or moved on. The specimen represents a novel microhabitat consisting of a remnant cuticle originally secreted by *Relicanthus daphneae*, itself a rarely encountered and recently described species, which occurs between 1,200 and 4,000 m, and microbial community living on and beneath its cuticle and tissue. These significant discoveries enabled by collection would have been unlikely were it not for its unusual golden color and the specimen’s mysterious egg-like appearance. In light of what we have learned from this event, we encourage the collection of more unusual specimens. Integrated analysis, a combination of detailed video imagery as well as specimen sampling and analysis are crucial for advancing our understanding of organismal biology and marine communities, possibly in unexpected and profound ways. We also highlight that the presentation herein offers the likely possibility of further questions and additional exploration of these poorly understood habitats. Finally, we note the proliferation of erroneous species occurrences showing up in important biodiversity resources, such as GBIF and OBIS, as a result of faulty identification of metabarcodes derived from environmental samples. Better quality control on sequence identifications going into these resources is warranted.

## Acknowledgements

The authors would like to extend an appreciation to the NOAA Ship *Okeanos Explorer* officers and crew, ROV operations team from the Global Foundation for Ocean Exploration, and NOAA Ocean Exploration and NOAA Research staff for the Golden Orb collection and the public outreach upon its discovery during expedition EX2306. Additionally, we are grateful to the science party of FK210605 (Randi Rotjan, Chief Scientist), and the Schmidt Ocean Institute, as well as R/V *Falkor* officers, crew, and pilots of the ROV SuBastian, who collected the central Pacific specimen with support from NOAA’s Deep Sea Coral Research and Technology Program in partnership with the University Corporation for Atmospheric Research (subaward # SUBAWD002484 to R. Rotjan) and S. Auscavitch who received postdoctoral support under a NOAA subaward 1305M322PNFFP0454 at Boston University. The authors also thank Katrin Linse (British Antarctic Survey) for facilitating comparative material and data of the *Relicanthus* specimen from the Southern Ocean; the collection of *R. daphneae* at the East Scotia Ridge was supported by the technicians of the ROV *Isis* during JC80, the JC80 scientific expedition crew, and funded the UK NERC Consortium Grant (NE/DO1249X/1) ChEsSo (PI: P. Tyler). Thanks to the Ocean Exploration Trust for imagery of *Relicanthus* anemones observed during expeditions NA035 and NA137. Initial laboratory work was supported in part by NMNH Postdoc Paula Rodríguez Flores who conducted the first initial cox1 barcodes of USNM 1699903. Isabel Pen provided UCE data for two of the actiniarian samples and Claudia Vaga was instrumental in tracking down key comparative sequences. Finally, we acknowledge the Smithsonian NMNH Laboratories of Analytical Biology (LAB) for resources associated with the generation of genetic data and scientific imaging at the National Museum of Natural History, Smithsonian Institution for equipment for imaging.

## Data Accessibility

NCBI BioSample records: SAMN56737688-694

NCBI SRA records: SRR37877116-122

NCBI Nucleotide records for mitochondrial genomes: PZ234282-83

## Supplementary Files

- Table S1. Small Subunit Ribosomal RNA assembled with PhyloFlash.
- Table S2. Samples included in UCE analysis.
- Table S3: Metagenomic assemblies associated with ‘Golden Orb’ samples.
- Table S4. Occurrence records of *Relicanthus daphneae*.
- Figure S1. Maximum likelihood phylogeny of Anthozoa estimated from a UCE dataset (94 loci) using a 50% occupancy matrix, with clades not collapsed. For information on samples see Table S2.

