## Supplementary Table S3 for "The Curious Case of the Golden Orb – Relict of *Relicanthus daphneae* (Cnidaria, Anthozoa, Hexacorallia), a deep sea anemone"

**Supplementary Table S3:** Metagenomic assemblies, GTDB taxonomy, and features associated with 'Golden Orb' samples.

| MAG ID (short) | Quality Tier* | GTDB taxonomy (to genus) | msa_percent | MAG Features |
|---|---|---|---|---|
| 1699903_Internal-deep.1 | HQ | Gammaproteobacteria; o__WLWR01; f__WLWR01; g__ | 96.9 | Near-complete MAG (117/120 markers) from a **deeply novel Gammaproteobacteria lineage**, likely a heterotroph in internal tissue. |
| 1699903_Internal-deep.8 | HQ/MQ | Gemmatimonadota; Longimicrobiales; f__UBA6960; g__ | 90.3 | High-quality **Gemmatimonadota** MAG from UBA6960, a poorly characterized sediment lineage; adds genomic detail on its role in deep-sea organic matter processing. |
| 1699903_Surface.1 | MQ–HQ | Chloroflexota; Dehalococcoidia; o__UBA2979; g__UBA6627 | 85.7 | Near-complete **Chloroflexota (Dehalococcoidia)** genome from the surface; likely involved in degradation of complex organics and redox cycling at the tissue–water interface. |
| 1699903_Surface.10 | MQ–HQ | Bacteroidota; Flavobacteriales; Flavobacteriaceae; g__CANLXL01 | 87.6 | Novel **Flavobacteriaceae** MAG associated with *Relicanthus* cuticle surface; Bacteroidota are classic polymer degraders, so this genome likely targets labile host-derived substrates. |
| 1699903_Surface.12 | MQ–HQ | Verrucomicrobiota; Chthoniobacterales; Terrimicrobiaceae; g__CAMASH01 | 87.1 | Verrucomicrobiota MAG from a **Terrimicrobiaceae-like lineage**, indicating colonization of the rock-hold surface by taxa often linked to complex carbon turnover. |
| 1699903_Surface.17 | MQ | Planctomycetota; Planctomycetia; Planctomycetales; Gimesia | 85.2 | **Planctomycetota (Gimesia)** MAG; planctomycetes frequently inhabit biofilms and surfaces, suggesting an important role in **surface-attached degradation** on the rock-hold. |
| 1699903_Surface.22 | HQ | Alphaproteobacteria; Rhizobiales; Rhizobiaceae; g__SZUA-100 | 96.0 | High-quality **Rhizobiaceae** MAG from the rock surface; represents a nearly complete Alphaproteobacteria genome that likely participates in **nitrogen and carbon cycling** on *Relicanthus* tissue. |
