## Supplementary figures and images for "The Curious Case of the Golden Orb – Relict of *Relicanthus daphneae* (Cnidaria, Anthozoa, Hexacorallia), a deep sea anemone"

### Supplementary Figure S1

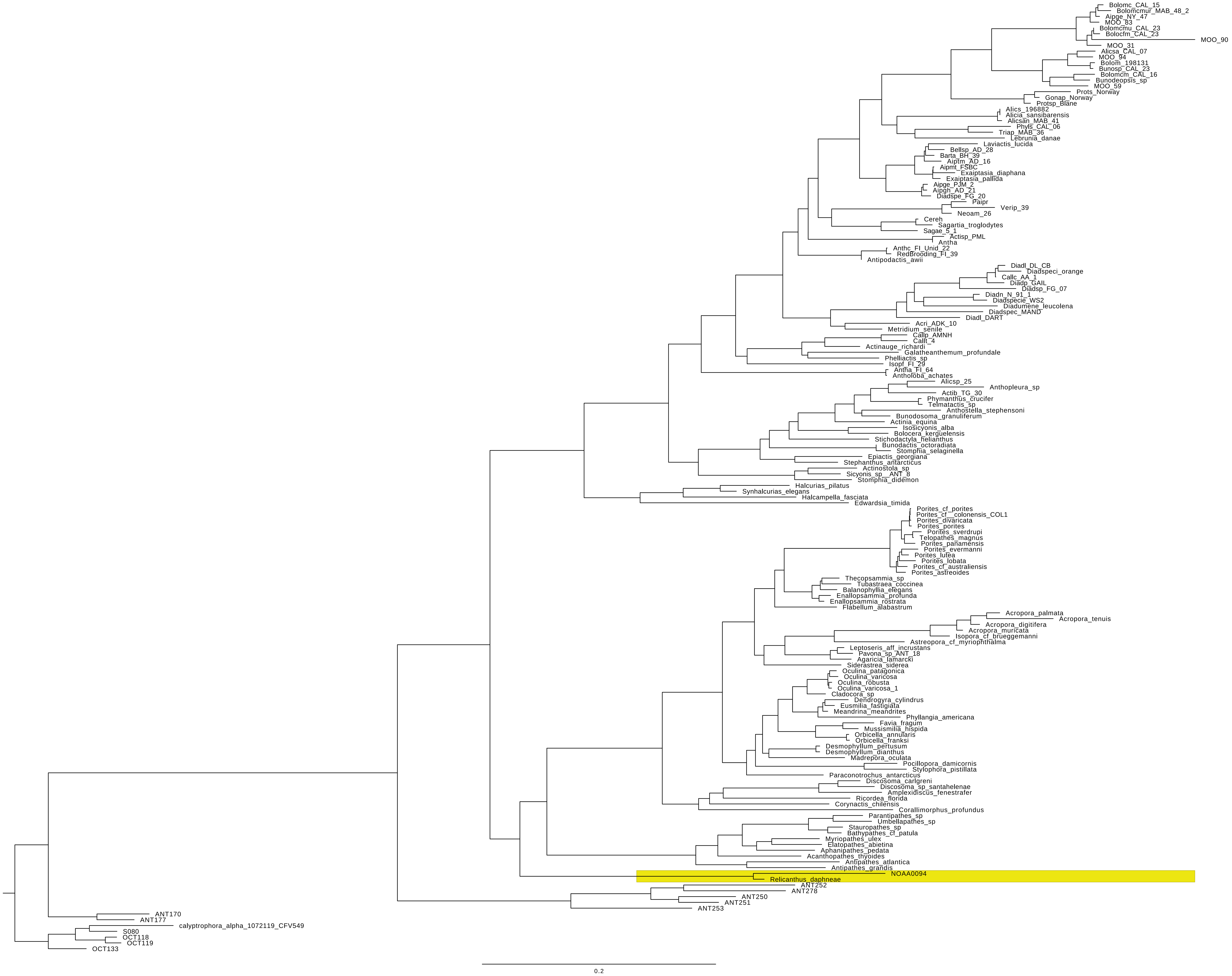
